# Speciation in *Heliconius* Butterflies: Minimal Contact Followed by Millions of Generations of Hybridisation

**DOI:** 10.1101/015800

**Authors:** Simon H. Martin, Anders Eriksson, Krzysztof M. Kozak, Andrea Manica, Chris D. Jiggins

**Author notes:** These authors contributed equally (Simon H. Martin, Anders Eriksson). Corresponding Author: S.H. Martin.

## Abstract

Documenting the full extent of gene flow during speciation poses a challenge, as species ranges change over time and current rates of hybridisation might not reflect historical trends. Theoretical work has emphasized the potential for speciation in the face of ongoing hybridisation, and the genetic mechanisms that might facilitate this process. However, elucidating how the rate of gene flow between species may have changed over time has proved difficult. Here we use Approximate Bayesian Computation (ABC) to fit a model of speciation between the Neotropical butterflies *Heliconius melpomene* and *Heliconius cydno*. These species are ecologically divergent, rarely hybridize and display female hybrid sterility. Nevertheless, previous genomic studies suggests pervasive gene flow between them, extending deep into their past, and potentially throughout the speciation process. By modelling the rates of gene flow during early and later stages of speciation, we find that these species have been hybridising for hundreds of thousands of years, but have not done so continuously since their initial divergence. Instead, it appears that gene flow was rare or absent for as long as a million years in the early stages of speciation. Therefore, by dissecting the timing of gene flow between these species, we are able to reject a scenario of purely sympatric speciation in the face of continuous gene flow. We suggest that the period of minimal contact early in speciation may have allowed for the accumulation of genomic changes that later enabled these species to remain distinct despite a dramatic increase in the rate of hybridisation.

## Introduction

Speciation is widely viewed as the development of reproductive isolation between lineages. However, there is now considerable evidence that reproductive isolation is not necessarily a genome-wide phenomenon, but rather that species integrity can be maintained despite gene flow affecting a considerable proportion of the genome [1–6]. What remains less clear is the importance of gene flow (or lack thereof) for the establishment of new species. Theory has shown that it may be possible, under certain genetic and selective conditions, for species to become established in the face of ongoing gene flow [7–14]. To test this theory, it is necessary either to observe speciation in real time, or to reconstruct the historical extent and timing of gene flow between existing species.

In geographic terms, speciation can be described as sympatric, parapatric or allopatric. We follow Mallet et al. [15] in defining these terms: sympatric populations share the same geographic area (but not necessarily the same niche), such that individuals from the two populations are liable to encounter one-another frequently over much of their range. Parapatric populations “occupy separate but adjoining geographic regions,” such that only a small fraction of individuals at the edge of each range are liable to encounter the other. Allopatric populations are geographically separated, such that encounters between them are very rare or impossible. Despite the abundance of closely related sympatric species, there are very few cases in which it can be stated with any certainty that speciation occurred in sympatry [e.g. crater-lake cichlids [16]]. In terms of gene flow, we can predict that sympatric speciation might involve a gradual decline over time, with higher rates of historic than contemporary gene flow [17]. In allopatric speciation, gene flow would be absent until the populations came into secondary contact. Parapatric speciation might fall somewhere in between these extremes, with a low level of gene flow throughout, potentially decreasing over time, but possibly also increasing if the populations later return to a sympatric distribution. Genomic data now offers the exciting possibility of reconstructing the history of gene flow between existing species, illuminating the roles of gene flow and geography in the origin of new species.

A number of methods exist to fit a model of “isolation with migration” (IM) using patterns of DNA sequence variation, thereby testing for post-speciation gene flow. This can be achieved by maximising the likelihood of observed genetic data in a coalescent framework, either directly [18–22] or using Markov Chain Monte Carlo (MCMC) approximation [23–26]. However, these approaches lack power and accuracy to examine change in the rate of gene flow over time [27,28], owing to characteristics of the standard IM model itself [29]. This limitation could be overcome through the implementation of more complex models [30], but this is currently not feasible in a likelihood framework. Approximate Bayesian Computation (ABC) offers a tractable means to fit such complex genetic models by avoiding the need to derive likelihoods [31]. ABC is therefore suited to the problem of reconstructing changes in the rate of gene flow during speciation [32,33], offering the potential to resolve long-standing debates in the speciation literature.

Here we investigate the history of gene flow during speciation in *Heliconius* butterflies. This Neotropical genus is well known for its broad diversity of aposematic wing patterns, and multiple instances of Müllerian mimicry – where unrelated species converge in wing pattern, providing a unified signal of toxicity to predators. Closely related species usually differ in wing pattern, and it is thought that pattern divergence between populations adapting to mimic different locally-abundant patterns could lead to parapatric speciation [34,35]. We examine *Heliconius melpomene* and *Heliconius cydno*, closely related species that have diverged in wing pattern and other ecological traits during the past million years, but continue to hybridise at low frequency [36] where their ranges overlap in the western parts of South America and Central America. The ability to compare sympatric and allopatric populations of *H. cydno* and *H. melpomene* provides an ideal opportunity to detect the genetic signatures of recent gene flow. Indeed, whole-genome studies have found evidence of abundant gene flow between these species, affecting a large proportion of the genome and extending deep into the past [3,4]. However, it has remained uncertain whether this pair diverged in the face of ongoing gene flow, or experienced a period of isolation early during speciation.

We used ABC to reconstruct the genealogical history of three populations: *H. cydno* from Panama, *H. melpomene rosina* from Panama (sympatric with *cydno*) and *H. melpomene melpomene* from French Guiana (allopatric) (Fig. 1A). Our model allowed for hybridisation between *H. cydno* and *H. melpomene* throughout speciation, and accounted for the possibility of a change in the rate of hybridisation during this time by considering two separate periods with distinct migration rates, the duration or which could vary. This enabled us to test various hypotheses under a single model, including a clean split without gene flow, continuous gene flow throughout speciation or gene flow restricted to ancient or recent time periods.

**Figure 1.**
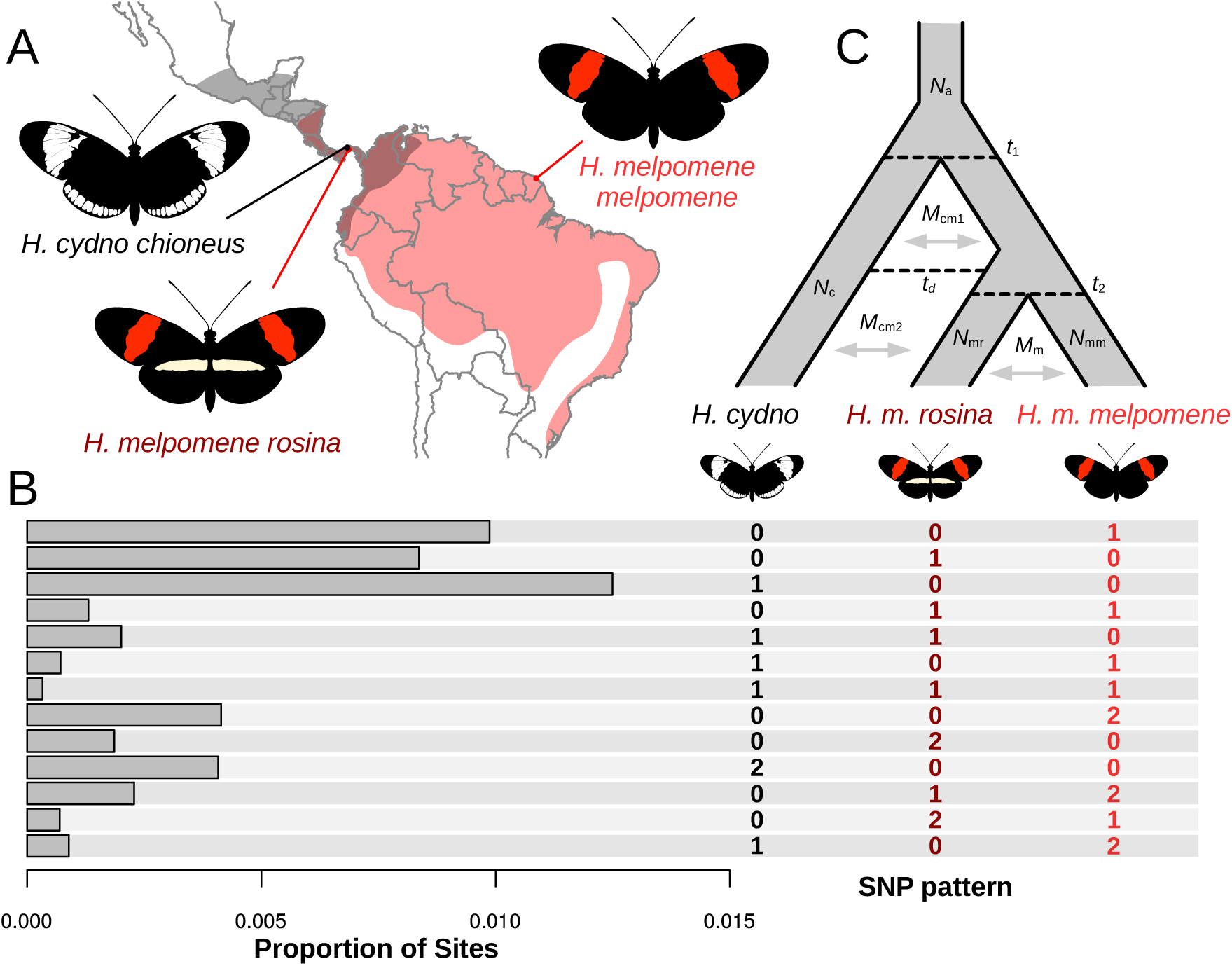
Species distributions, sample locations, summary statistic and model design. **A.** Distributions (shaded) of *H. melpomene* (light red) and *H. cydno* (grey), based on Rosser et al. [37]. Sampling locations in Panama and French Guiana are indicated. **B.** The composite summary statistic used consisted of the proportions of the 13 possible biallelic genotype combinations, where one individual is sampled from each population (given to the right). ‘0’ and ‘2’ indicate alternative homozygous states and ‘1’ indicates the heterozygous state. Given that four individuals were sampled from each population, the proportion of biallelic SNPs carrying each pattern was averaged over all 64 possible sets of three samples. Although 25 SNP states are theoretically possible, twelve of these can be folded if we ignore major and minor alleles (e.g. 2-0-1 is equivalent to 0-2-1) and so these were counted together, to give 13 unique states. **C.** The model had ten free parameters: four population sizes (*N*_a_, *N*_c_, *N*_mr_ and *N*_mm_) (the ancestral *H. melpomene* population size was assumed to be the average of *N*_mr_ and *N*_mm_); migration rates between *H. cydno* and *H. melpomene* in Periods 1 and 2 (*M*_cm1_ and *M*_cm2_) the time dividing Periods 1 and 2 (*t*_d_); migration rate between the two *H. melpomene* populations (*M*_m_), and the split times for the two species (*t*_1_) and the two *H. melpomene* populations (*t*_2_).

## Results

### Genotype data and summary statistics

Whole genome resequence data from twelve wild-caught butterflies, with four representatives from each of the three sampled populations, *H. cydno*, *H. m. rosina* and *H. m. melpomene*, were used for model fitting (S1 Table, data from Martin et al. 2013). Only intergenic regions, as defined by the *Heliconius melpomene* reference genome annotation v1.1 [38], with high-quality genotype calls (see Materials and Methods) for all twelve samples, were considered. We also excluded all scaffolds on the Z chromosome, which is known to experience strongly reduced gene flow [4], as well as putative CpG clusters, which can have unusual mutation rates. These criteria gave ~60 million sites (22% of the genome), of which approximately 10% were polymorphic (Single Nucleotide Polymorphisms, SNPs). The composite summary statistic used for model fitting consisted of the proportion of bi-allelic sites carrying each possible combination of genotypes among three diploid individuals. These proportions were averaged over all possible triplets, where each population is represented by one individual, and folded such that major and minor alleles were not distinguished (Fig. 1B, see Materials and Methods for details). This composite summary statistic is similar to a three-dimensional site frequency spectrum. It provides a nearly exhaustive summary of the available SNP data among the ingroup taxa, is independent of linkage effects and scalable to any number of sites.

As expected, the most common SNP patterns were singletons, where one individual was heterozygous and the other two were homozygous for the same allele (0-0-1, 0-1-0, and 1-0-0; Fig. 1B). The most common pattern overall was 1-0-0, where *H. cydno* is heterozygous and both *H. melpomene* individuals are homozygous. This is unsurprising, given the longer branch leading to *H. cydno* (Fig. 1B). The pattern 0-0-1, where *H. m. melpomene* from French Guiana is heterozygous, was also considerably more common than 0-1-0, where *H. m. rosina* is heterozygous. This is consistent with increased shared variation between *H. cydno* and the sympatric *H. m. rosina.* Similarly, 1-1-0 was more common than 1-0-1, and 0-0-2 was more common than 0-2-0.

### Estimating the timing and extent of gene flow

We consider a model with three populations (Fig. 1C), corresponding to *H. cydno* (which splits from the ancestral *melpomene* population at time *t*_1_), and *H. m. rosina* and *H. m. melpomene* (which split at time *t*_2_). Each lineage has a separate population size, except for the ancestral *melpomene* population, which is assumed to have a size equal to the mean of the two *melpomene* populations. Because the two *melpomene* populations represent extremes of a somewhat continuous range, migration between them is allowed at a continuous rate *M*_m_. Migration is also allowed between *H. cydno* and *H. melpomene*, although after the split between the *melpomene* populations (*t*_2_) only *H. m. rosina* is able to exchange migrants with *H. cydno.* Two distinct periods of between-species migration are modelled, with rates *M*_cm1_ and *M*_cm2_. These periods are divided at a time *t*_d_ such that Period 1 begins at *t*_1_ and ends at *t*_d_, and Period 2 runs from *t*_d_ to the present. This model had ten free parameters: four population sizes, two split times, three migration rates and one time dividing the migration periods. Model parameters were estimated using Approximate Bayesian Computation (ABC) based on the summary statistics described above (see Materials and Methods for details). Uniform priors were used for all parameters except for *t*_1_ and *t*_2_, for which prior distributions were estimated by analysis of mitochondrial sequence data (see Materials and Methods for details).

Our model pointed toward a dramatic change in the rate of inter-specific migration (i.e. hybridisation resulting in gene flow) from early to later stages of speciation (Fig. 2). Migration was minimal in Period 1 (*M*_cm1_~0.08 migrants per year [posterior mean]), and around tenfold greater in Period 2 (*M*_cm2_~0.81 migrants per year) (Fig. 2A, Table 1). The date of transition between these two periods (*t*_d_) had a fairly wide posterior distribution, with a mean of 0.5 million years ago (Ma), but a 90% posterior density interval extending from 0.1 Ma to 1.2 Ma (Fig. 2A, Table 1). This was nevertheless considerably more recent than the inferred time of speciation (*t*_1_), which had a fairly narrow posterior distribution centred around 1.5 Ma (Fig. 2A, Table 1). Therefore, our results support a case of stronger isolation during the early stages of speciation, with a large increase in the rate of hybridisation later. The posterior mean of 0.5 Ma for the date dividing the two periods would imply that hybridisation was rare or absent during the first two thirds of the time since initial divergence (~1 million years). However, we note that this transition cannot be dated with great accuracy, and indeed our model does not allow us to infer whether the increase in hybridisation was sharp or gradual. Nevertheless, it is notable that the posterior mean for the onset of more frequent hybridisation coincides roughly with the split between the two *H. melpomene* populations (*t*_2_~0.53 Ma, Fig. 2A). Interestingly, the inferred rate of gene flow between *H. cydno* and *H. m. rosina* during this second period is several times greater than that between the two *H. melpomene* populations (Fig. 2A, Table 1).

**Figure 2.**
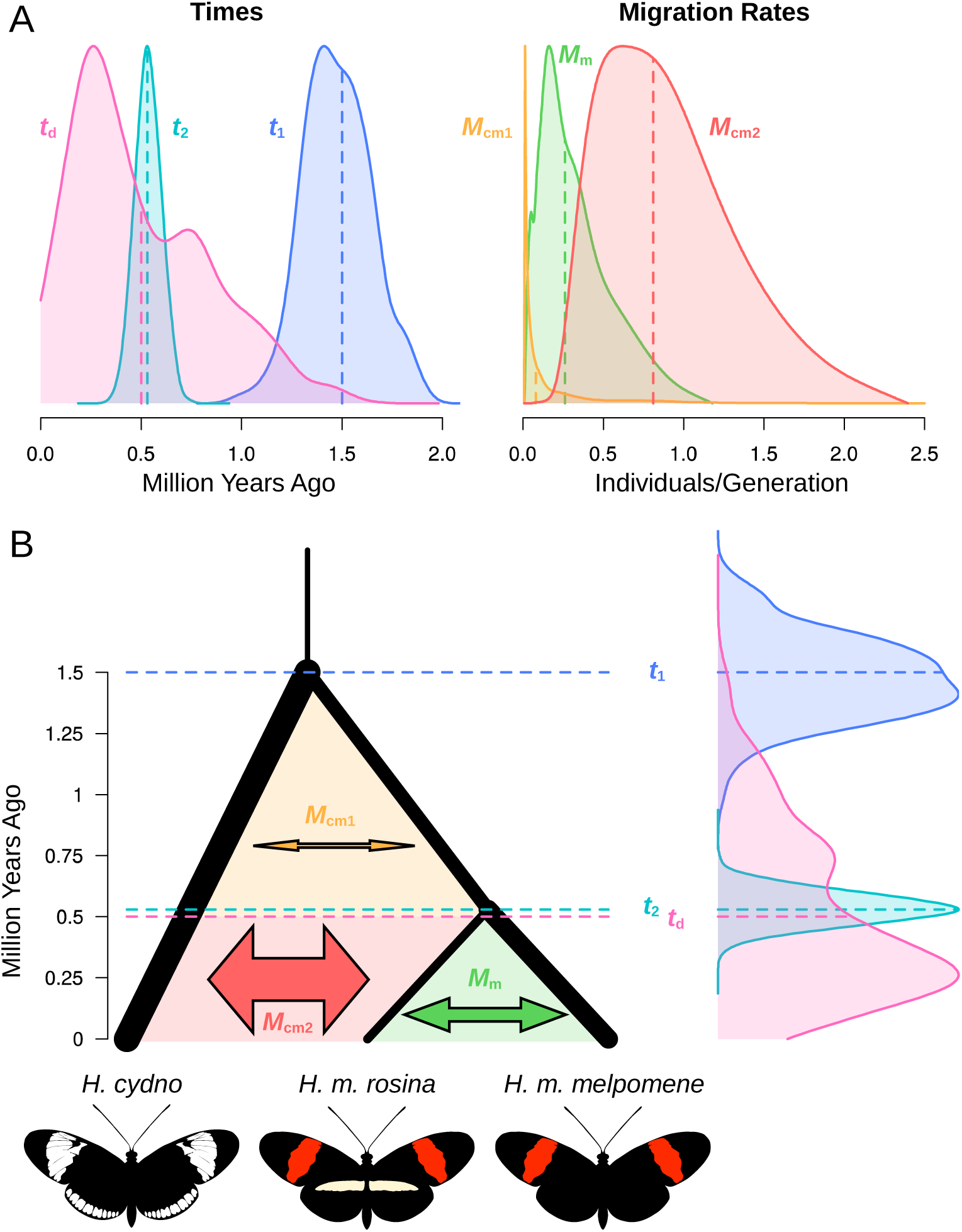
Posteriors for time and migration rate parameters, and a schematic representation of the inferred model. **A.** Posterior distributions for the three time perameters (left) and three migration rates (right). Densities are scaled for ease of comparison. See S2 Figure for all posterior distributions. Posterior means are indicated by vertical dashed lines. **B.** A schematic phylogeny, where the posterior mean for each parameter is indicated. Colours correspond to those in A. Times are indicated by horizontal dashed lines. Posteriors for the times are given on the right-hand-side for reference. Migration rates are indicated by arrows, with the width of the arrow scaled according to the migration rate. Relative population sizes are indicated by branch widths. See Table 1 for all values.

**Table 1.** Properties of posterior distributions for the ten model parameters

| Parameter | Mean | Mode | Median | HPD90<br>Lower<sup>1</sup> | HPD90<br>Upper<sup>1</sup> |
| --- | --- | --- | --- | --- | --- |
| $t_1$ (Ma) | 1.5 | 1.4 | 1.5 | 1.2 | 1.8 |
| $t_2$ (Ma) | 0.53 | 0.53 | 0.53 | 0.42 | 0.65 |
| $t_d$ (Ma) | 0.5 | 0.3 | 0.4 | 0.1 | 1.2 |
| $M_{cm1}$ (ind. /year) | 0.08 | 0.01 | 0.06 | 0.01 | 0.86 |
| $M_{cm2}$ (ind. /year) | 0.81 | 0.95 | 0.84 | 0.36 | 1.6 |
| $M_m$ (ind. /Year) | 0.26 | 0.34 | 0.28 | 0.06 | 0.77 |
| $N_c$ (M ind.) | 5.3 | 5.3 | 5.3 | 4.6 | 5.9 |
| $N_{mr}$ (M ind.) | 1.8 | 1.8 | 1.8 | 1.2 | 2.4 |
| $N_{mm}$ (M ind.) | 3.9 | 3.9 | 3.9 | 3.4 | 4.4 |
| $N_A$ (M ind.) | 1.1 | 1.3 | 1.1 | 0.3 | 2.0 |

<sup>1</sup> Upper and lower bounds of the 90% Highest Posterior Density (HPD) interval. The HPD is defined as the set of values making up 90% of the density distribution and within which all values have higher density than outside.

Estimates of *N_e_* were generally larger than previous estimates for these species [3], with relatively narrow posterior distributions (Table 1, S2 Figure). This difference may be driven by a lower mutation rate used in the present study. Consistent with this previous study, the estimated *N_e_* for *H. cydno* (~5.3 M individuals) was larger than both *H. m. rosina* (~1.8 M) and *H. m. melpomene* (~3.9 M).

Overall, simulated summary statistics from the retained parameter combinations matched the observed summary statistic well (S3 Figure). To investigate the robustness of our conclusions, we repeated the ABC analysis using the split time priors from model F (S3 Table and see above). The general findings were largely the same (data not shown), indicating that our conclusions are not strongly influenced by the split time priors.

## Discussion

The existence of distinct species that share genetic material through hybridisation, or have done so in the recent past, is no longer disputable. Genome-scale data have provided overwhelming evidence of pervasive gene flow between species, in taxa as diverse as fruit flies [39], flycatchers [2], and hominids [19,40]. However, understanding the timing of gene flow during speciation, and the importance of geographical isolation for species establishment remain difficult. Here, we combine a large-scale genomic dataset with Approximate Bayesian Computation (ABC) to reconstruct speciation in *Heliconius* butterflies. Our findings support recent studies showing abundant gene flow between *H. melpomene* and *H. cydno* [3,4], and suggest that gene flow has been ongoing for approximately half a million years (two million generations). However, we also find that hybridisation was rare or absent during the roughly the first million years of divergence between these species, a factor that may have played an important role in their establishment.

In order to reduce the parameter space explored, we used a fairly narrow joint prior distribution for the two split times, *t*_1_ and *t*_2_. This joint distribution was inferred using mitochondrial sequence data, under the assumption that mitochondrial introgression between *H. cydno* and *H. melpomene* should be unlikely. This is reasonable given the fact that female hybrids are sterile [41]. Although one *H. melpomene* population from Colombia is known to carry *H. cydno-like* mitochondrial haplotypes [42], an analysis of all 449 unique haplotypes available on Genbank has not indicated any other mitochondrial introgression events between these species (see Materials and Methods). Our *cydno-melpomene* split time of 1.5 Ma is within the range of values inferred in previous analyses using IM-based methods [3,42,43]. In addition, posterior distributions for both split times were not strongly skewed toward the edge of the priors. It is nevertheless possible that the mitochondrial split times provide inadequate estimates of the nuclear split times. Even if this were the case, it is likely that our general conclusion of reduced migration earlier in speciation will still hold. Indeed, when we repeated the ABC analysis with a different, but overlapping, set of split time priors, the results were largely unchanged.

Allopatric speciation is often the null hypothesis in speciation studies [44]. One scenario that would be consistent with our results is that speciation began in allopatry, possibly with the emerging species separated by the Andes mountains. Subsequent range expansion of *H. melpomene* into the western Andes and Central America would have lead leading to secondary contact. The lower *N_e_* of *H. m. rosina* compared to *H. m. melpomene* is consistent with range expansion from east to west, which has been proposed previously [45]. However, our results may also be consistent with parapatric speciation, which is probably more common in *Heliconius* [35,46]. In fact, most pairs of sister species in the genus are sympatric or parapatric (Rosser et al. In Review). Allopatric populations of extant species, for example those on Caribbean islands, tend not to display phenotypic and ecological divergence from their mainland progenitors. In contrast, many species, including *H. melpomene* and *H. cydno,* are divided into numerous parapatric wing-pattern races across their mainland ranges. *H. melpomene* and *H. cydno* are also partially segregated by altitude, so it is plausible that parapatric adaptation to altitude in the Andes played a role in their speciation. The evolution of strong assortative mating associated with wing pattern might then have led to nearly complete reproductive isolation between the parapatric populations. Indeed, loci affecting both mate preference and hybrid sterility are known to be physically linked to wing patterning loci in these species, which might have enhanced reproductive isolation following divergence in wing pattern [47,48]. Ecological divergence, most notably in host plant use but perhaps also microhabitat preference [49], would then have followed later, permitting sympatric coexistence without competition [34]. The inferred increase in gene flow later in speciation might therefore reflect increased contact associated with the transition from parapatric to sympatric ranges. One final piece of evidence for parapatric speciation is the existence of several other species pairs that may represent an intermediate step in this process. The best studied are *Heliconius himera* and *H. erato*, which are largely parapatric with only narrow zones of overlap. They are strongly differentiated genetically [50] and display assortative mating based on colour pattern [51]. However, they have not diverged in host plant usage [52], which perhaps prevents their sympatric coexistance through competitive exclusion.

Regardless of its cause, we can speculate that an initial period of reduced gene flow contributed to the formation of these species. Reduced gene flow can facilitate the accumulation of Dobzhanzky-Muller incompatibilities [53,54], which would help to maintain species integrity even after the rate of hybridisation increased. For example, gene flow is minimal across the entire Z chromosome [4], consistent with a high density of incompatibility loci in this part of the genome. Interestingly, in these species there are also genetic associations between wing pattern, host preference and mate preference loci which likely facilitate coexistence in sympatry [47,48]. However it is unclear whether such associations have arisen since hybridisation became widespread, or whether they fortuitously pre-dated the period of extensive contact. Finer-scale analysis of the patterns of introgression across the genome, combined with mapping incompatibility loci and structural differences in the genome will help to dissect the various factors contributing to species persistence.

Despite our finding that hybridisation was rare or absent for approximately two thirds of the time since speciation, this nevertheless implies that the hybridisation has now been ongoing for around two million generations. Our model assumes a single change in the rate of gene flow, while it is highly probably that this rate has changed more gradually through time. The wide posterior distribution for the time of this transition is consistent with a gradual increase over perhaps hundreds of thousands of generations. There is also reason to believe that the rate of gene flow has recently begun to decrease. The existence of character displacement (sympatric males display stronger mate discrimination than allopatric males [55], suggests that selection may have acted to reinforce reproductive isolation in sympatry. Nevertheless, there remains strong evidence that gene flow continues today, both in the occurance of natural F1 hybrids [36], and in geographic patterns of shared variation [4]. Specifically, *H. cydno* samples from Panama share an excess of variation with *H. melpomene* samples from the same location compared to those from about 100 km away [4]. A recent study of the hybridising mouse subspecies *Mus musculus musculus* and *M. m. domestica* supported a similar scenario to that described here, with gene flow occurring over the last 25% of time since initial divergence, although at a lower rate (<0.2 ind/gen) [32]. Our estimate of ~0.84 migrants per generation represents the effective number of hybrids, but it is certain that the number of actual F1 hybrid butterflies produced exceeds this value considerably, for two reasons. Firstly, in accordance with Haldane’s Rule, female (ZW) F1 hybrids are sterile [41], and therefore do not contribute to observed gene flow. Secondly, F1 hybrids are subject to increased predation owing to their non-mimetic wing patterns [56].

Finally, it is worth considering the consequences of continued hybridisation between these species. Although a whole-genome phylogeny groups the *H. melpomene* populations as monophyletic, currently 40% of 100 kb windows group *H. m. rosina* with *H. cydno,* to the exclusion of the French Guianan *H. m. melpomene* [4]. Nevertheless, these sympatric populations retain the phenotypic, behavioural and ecological traits specific to their respective species, implying that species integrity is surprisingly resilient to gene exchange. It is certain that gene flow is inhibited by selection in some genomic regions, most obviously the wing pattern loci. However, natural selection has also favoured the occasional exchange of wing pattern alleles between certain populations of these clades, producing the paired mimetic races of *H. melpomene* and *H. timareta* found on the eastern slopes of the Andes [38,57]. It seems likely that much of the genome is neutral with respect to gene flow, and that most of the signal seen here is due to neutral exchange of alleles in sympatry, although we have not attempted to test for evidence of adaptive introgression. It is therefore possible that ongoing hybridisation, even at a low rate, might eventually lead to a situation where the majority of the genome clusters populations by geography rather than by species, making one or both species paraphyletic. It seems inevitable that genomic studies will reveal such species pairs in the near future, posing a challenge to species definitions based on aggregate genetic ancestry.

## Materials and Methods

### Samples and genotyping

We used published whole-genome resequence data for twelve wild-caught butterflies (S1 Table, data from Martin et al. [4], www.datadryad.com doi:10.5061/dryad.dk712). Details of the sequencing, mapping and genotyping procedures are described by Martin et al. [4]. Briefly, 100bp paired-end Illumina reads were mapped to the *H. melpomene* reference genome [38], version 1.1, using Stampy [58]. Local realignment around indels and genotyping were both performed using The Genome Analysis Toolkit (GATK) [59]. For the purpose of this study, we considered only intergenic SNPs, identified based on the *H. melpomene* genome annotation, version 1.1. CpG islands were identified using the program CpGcluster [60], and these sites were excluded. Only high quality genotype calls were considered. High quality genotypes met the following conditions: quality (QUAL) ≥ 30, 10 ≤ read depth per individual ≤ 200, and GQ ≥ 30 for SNPs. Processed genotype calls data are available from www.datadryad.com doi:XXX.

### Summary statistic

The summary statistic used for model fitting was a composite of the proportion of sites representing each of the possible combinations of bi-allelic genotypes among three diploid individuals, with one individual representing each population (Fig. 1B). For example, a SNP would be assigned the pattern 0-1-2, if the *H. cydno* individual was homozygous, carrying zero copies of the minor allele, the *H. m. rosina* individual was heterozygous, carrying one copy, and the *H. m. melpomene* individual was homozygous with two copies of the minor allele. The counts of all patterns were then folded, such that major and minor alleles were not taken into account. For example the pattern 0-1-2 was taken as equivalent to 2-1-0. This gave 13 unique SNP patterns (Fig. 1B). Because four individuals were sampled from each population, the counts of each pattern were averaged over all 64 possible triplets with one individual from each population. Custom scripts used to calculate and plot summary statistics are avauialable from www.datadryad.com doi:XXX. Since having too many summary statistics is a known problem with ABC, we used Partial Least Squares [implemented in the findPLS.r script in the ABC Toolbox [61]] to find the eight most informative linear combinations of the original summary statistics.

### Model

A three-population model of isolation with migration was used (Fig. 1C). An ancestral population divides at time *t*_1_ into two populations (corresponding to *H. cydno* and *H melpomene*, respectively). At time *t*_2_, the *melpomene* population further divides into the *H. m. rosina* and *H. m. melpomene* races, which remain connected by limited gene flow at a constant rate *M*_m_. The two *H. melpomene* populations, the *H. cydno* population, and the ancestral population, all have unique population sizes, but the size of the ancestral *melpomene* population is assumed to be the average of the two *melpomene* populations. Migration is allowed between *H. cydno* and the ancestral *H. melpomene* population, and between *H. cydno* and *H. m. rosina* after the two *H. melpomene* populations diverge. Two distinct periods of hybridisation are modelled, with rates *M*_cm1_ and *M*_cm2_. These two periods occupy the entire speciation time from *t*_1_ to the present, and are divided at time *t*_d_. Hence, Period 1 runs from *t*_1_ to *t*_d_ and Period 2 from *t*_d_ to the present. The division between the periods, *t*_d_, may fall anywhere between *t*_1_ and the present. A constant mutation rate of 1.9×10^−9^ per site per generation was used. This corresponds to the estimated per-generation mutation rate for *H. melpomene* [62], corrected for weak purifying selection on intergenic regions by multiplying by the relative level of interspecific divergence at intergenic and putatively neutral four-fold degenerate sites (data not shown). A generation time of 0.25 years was assumed [63].

### Priors for split times

To reduce the dimensionality of the model, we used fairly narrow priors for the two split times *t*_1_ and *t*_2_ (S2 Table). These were inferred using analysis of mitochondrial sequence data, which should be resistant to gene flow between these taxa. This assumption was first tested with a Maximum Likelihood analysis of all 847 publicly available sequences (449 unique haplotypes) in RAxML v.8 [64]. This identified a single, previously known [42] case of mitochondrial introgression between these species, which does not involve the populations considered here. Sequence data for 1606 bp of *CoI/II* for 125 samples from several populations of *H. melpomene*, *H. cydno* and the outgroup silvaniform clade were obtained from Genbank and the data of from Martin et al. [4], and aligned with MUSCLE [65]. Strict and relaxed molecular clock models and codon-partitioning schemes were fitted to the data in BEAST v. 1.8. [66] and compared with a posterior analog of AIC (AICM) and Bayes Factors calculated by the Stepping Stone Analysis [67] (S2 and S3 Table). For root-calibrated analyses, the split time between the *H. melpomene* and Silvaniform caldes inferred by Kozak et al. [68] was used. For fixed rate analyses a mutation rate of 0.0024 per million years was used, as inferred under a relaxed-clock model applied to the complete *Heliconiini* alignment of Kozak et al. [68]. While the exact split dates varied, all approaches converged on the same topology and a similar ratio of split times *t*_1_/*t*_2_ (S3 Table). Bayes Factors and AICM [67] favoured a strict clock model (model E, S2 Table) with a separate partition for third codon positions. We present results obtained using the split times from model E, although we also tested the split times from model F (S3 Table). The resulting joint posterior distributions for the two split times formed the priors for the ABC simulations, with pairs of times for *t*_1_ and *t*_2_ being drawn together from this joint distribution.

### Model fitting using ABC

Approximate Bayesian Computation (ABC) was used to estimate parameters of the model. Briefly, ABC fits a model by evaluating the distance between observed and simulated summary statistics, allowing the estimation of posterior probability distributions without calculating likelihood functions.

Uniform priors were used for all parameters except for the split times *t*_1_ and *t*_2_ (see above). Two million parameter combinations were generated over the parameter space by sampling from the prior distributions randomly and independently (except for parameters *t*_1_ and *t*_2_, which had a joint prior distribution). A custom program (written in C/C++) was then used to simulate 100,000 unlinked SNPs from our model under the standard coalescent framework [69] for each sampled parameter set, and then to calculate the summary statistic. SNPs were simulated independently (i.e. unlinked) as the composite summary statistic used here, which is based on the genome-wide joint frequency spectrum, should not be strongly influenced by linkage disequilibrium, especially given the fairly rapid decline in LD in *Heliconius melpomene* [4]. Using the standard ABC method, we used the Euclidian distance between simulated and observed values to identify parameter combinations that fit the data well. We used a cut-off of 0.01 for accepting parameter combinations, yielding 27377 good parameter combinations for ABC. To account for variation in goodness of fit obtained among retained parameters, the distribution of retained parameters was adjusted using the General Linear Model method of [70], as implemented by ABCestimator of the ABC Toolbox [61].

## Acknowledgements

We thank Nick Barton, Aylwyn Scally and Konrad Lohse, for helpful comments on earlier drafts of the manuscript. Computational support was provided by Jenny Barna, School of Life Sciences, University of Cambridge.

## List of supplemental files

**S1 Table. Sample information and sequencing read depth**

**S2 Table. Model comparison of various strategies for estimating the divergence times in BEAST.** Models E and F are equally good based on Bayes Factors. Log Bayes Factors (BF) are calculated based on the log Marginal Likelihood estimates (MLE) from Path Sampling (PS) and Stepping Stone Analysis (SSA) (Baele et al. 2012). The molecular clock rates were calibrated by either modelling the age of the root or setting a constant rate of substitution. UCLD= Uncorrelated Lognormal clock.

**S3 Table. Parameter values for the three best Bayesian models of divergence between the *CoI/II* sequences.**

**S1 Figure. Bayesian phylogeny of the *Cytochrome Oxidase I/II* (*CoI/II*) haplotypes from the *Heliconius melpomene* / *H. cydno* clade**. This tree was estimated under the relaxed molecular clock model (F) with calibrated age of the root. The sampling is based on a balanced design with 25 samples per group, including *H. timareta* as the sister species of *H. cydno*. Blue: *H. melpomene* from Central America; green: *H. melpomene* from French Guiana (clade including allopatric samples used in the analysis of the nuclear genomes); pink: *H. cydno*; grey: Silvaniform outgroups. Time scale in millions of years, bars represent the 95% HPD intervals around split ages. Samples with whole mitogenome data from Martin et al. (2013) indicated in capital letters. NCBI GI numbers are provided for all sequences obtained from GenBank.

**S2 Figure. Posterior density plots inferred by ABC for the 10 model parameters.** Parameters labels match Fig. 1 in the main text, with the three time parameters given in the left-hand column (A-C), migration rates in the middle column (D-F), and population sizes in the right-hand column (G-J). Grey shading indicates the 90% highest posterior density (HPD) interval, and posterior means are indicated by vertical dashed lines.

**S3 Figure. Summary statistics from retained parameter combinations compared to the observed values.** Patterns 1-13 correspond respectively to 0-0-1, 0-0-2, 0-1-0, 0-1-1, 0-1-2, 0-2-0, 0-2-1, 2-0-0, 1-0-0, 1-0-1, 1-0-2, 1-1-0 and 1-1-1. Box plots indicate the distribution of simulated values from the 27377 retained parameter combinations. Red points indicate outliers (outside of the 10th and 90th percentiles). Yellow diamonds indicate observed frequencies, as in Fig. 1B. Here the y-axis indicates the the scaled frequency of each pattern, in units of million generations. This can be obtained by dividing the observed absolute frequency of each pattern (as in Fig. 1B) by the mutation rate per site per million years.

